# Root electrotropism in Arabidopsis does not depend on auxin distribution but requires cytokinin biosynthesis

**DOI:** 10.1101/2020.07.30.228379

**Authors:** Maddalena Salvalaio, Nicholas Oliver, Deniz Tiknaz, Maximillian Schwarze, Nicolas Kral, Soo-Jeong Kim, Giovanni Sena

## Abstract

An efficient foraging strategy for plant roots relies on the ability to sense multiple physical and chemical cues in soil and to reorient growth accordingly (tropism). Root tropisms range from sensing gravity (gravitropism), light (phototropism), water (hydrotropism), touch (thigmotropism) and more. Electrotropism, also known as galvanotropism, is the phenomenon of aligning growth with external electric fields and currents. Although observed in a few species since the end of the 19^th^ century, the molecular and physical mechanism of root electrotropism remains elusive, limiting the comparison to more defined sensing pathways in plants.

Here we provide a first quantitative and molecular characterisation of root electrotropism in the model system *Arabidopsis thaliana*, showing that it does not depend on an asymmetric distribution of the plant hormone auxin, but that instead it requires the biosynthesis of a second hormone, cytokinin. We also show that the dose-response kinetics of the early steps of root electrotropism follows a power law analogous to the one observed in some physiological reactions in animals.

A future full molecular and quantitative characterisation of root electrotropism would represent a step forward towards a better understanding of signal integration in plants, and an independent outgroup for comparative analysis of electroreception in animals and fungi.

## INTRODUCTION

Plant roots navigate the complex soil environment in search of water and nutrients, through various sensing mechanisms reorienting their growth towards or away from signal sources (tropism) (Muthert et al., 2020). For example, hydrotropism redirects growth towards high moisture (Miyazawa and Takahashi, 2019), (negative) phototropism redirects away from light sources (Kutschera and Briggs, 2012), and gravitropism induces growth downwards, following the gravity vector (Su et al., 2017). The integration of overlapping and frequently contradicting molecular and physical signals is as critical for plant roots as it is for other soil organisms.

Although many molecular aspects of a few root tropisms have been understood (Muthert et al., 2020), several key sensing mechanisms remain elusive. One of these is the capacity of plant roots to sense electric fields (electrotropism or galvanotropism) (Navez, 1927). The local physical and chemical properties of soil determine the presence of mobile electrical charges and the generation of spontaneous electric fields (Pozdnyakov and Pozdnyakova, 2002). For example, electrostatic fields can appear from charge separation in minerals like clay (Ward, 1990), by electrokinetic conversion caused by a conducting fluid like water flowing through rocks (Revil et al., 2003) or by local accumulation of mineral ions important for plant metabolism such as ammonium and nitrate ions. Local electric fields in soil can have biological origin as well, for example from negatively charged bacteria (Olitzki, 1932) or from ions and charged molecules released by microorganisms (Chabert et al., 2015) or plant roots (Takamura, 2006). All this suggests that transient electrostatic fields in soil encode unique information regarding the localisation of water, micronutrients and organisms, and it is plausible that a sensing mechanism to detect such signal provides a selective advantage.

In fact, the reception of electric fields (electroreception) has been observed in vertebrates (Crampton, 2019) and invertebrates (Clarke et al., 2013), including common model systems such as *C. elegans* (Sukul and Croll, 1978) and *Dyctostylium* (Shanley et al., 2006), as well as in fungi (McGillivray and Gow, 1986). Interestingly, electric sensing structures have been identified only in a few aquatic animal species (Peters et al., 2007) and more recently in bumble-bees (Sutton et al., 2016).

First recorded in 1882 (Elfving, 1882) and rediscovered at the start of the 20^th^ century (Ewart and Bayliss, 1905), root electrotropism has been studied sporadically in maize (*Zea mays*), peas (*Pisum sativum*) and bean (*Vigna mungo*) but with contradicting results (Wolverton et al., 2000). Crucially, the anatomical and molecular details of sensing electric fields are still largely unknown in roots. A quantitative description of root electrotropism’s kinetics is also missing, preventing a comparative analysis with animal electroreception and electrotaxis.

## RESULTS

### A novel root electrotropism assay

To study root electrotropism in the plant model system *Arabidopsis thaliana*, we developed a new setup to stimulate, image and quantitate electrotropism in its primary root. Briefly, roots were grown vertically in a transparent chamber (V-box) containing a buffered liquid medium and two immersed electrodes connected to a power supply to generate a uniform electric field and an ionic current perpendicular to the growing roots (Fig. 1a and Methods).

**Figure 1.**
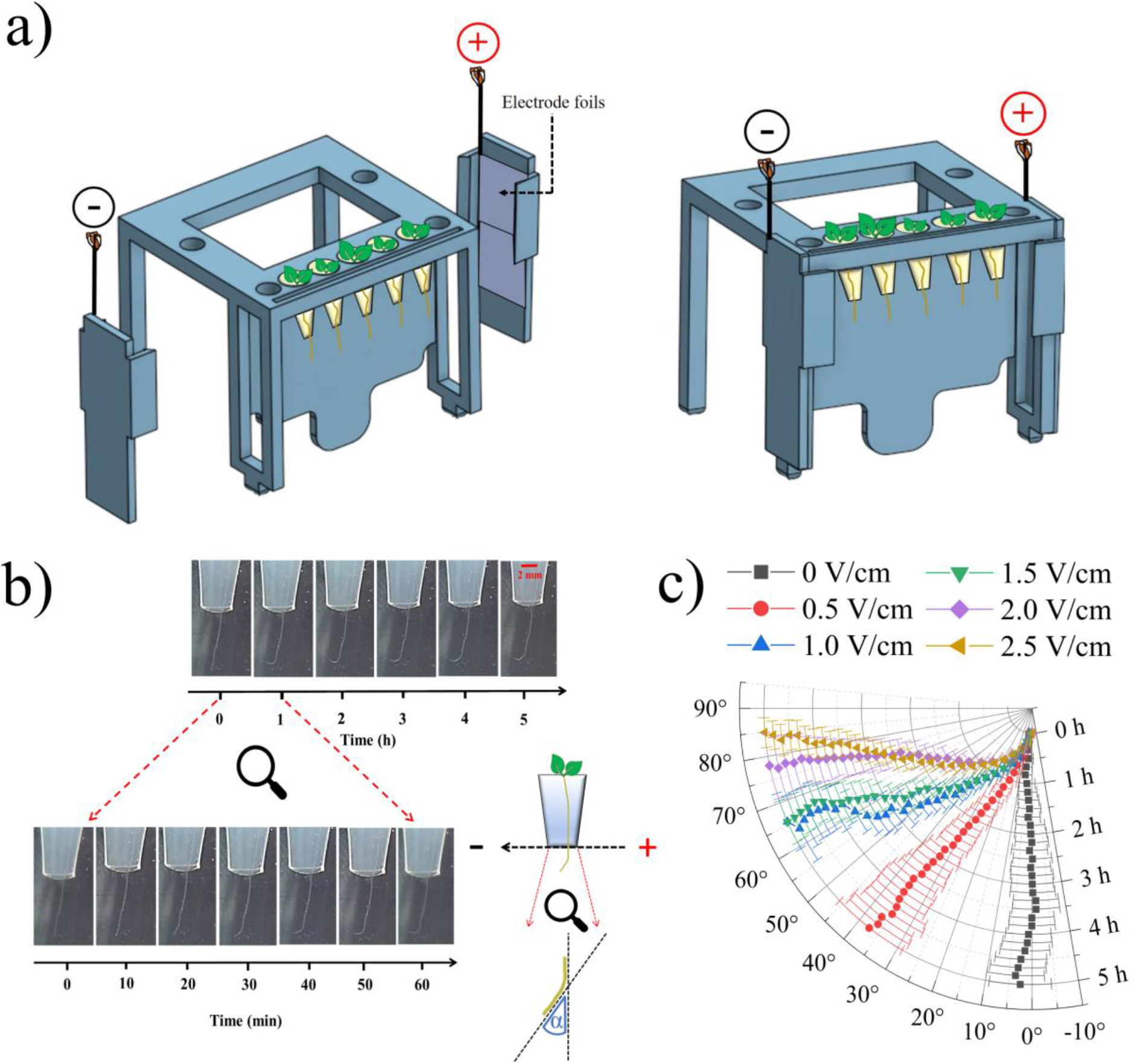
**(a)** Schematic of the 3D-printed module used in the V-box. **(b)** Representative time-lapse series of a single root as imaged by the Raspberry camera from the V-box; inset, schematic of the angle measured. **(c)** Polar plot of the average root tip orientations relative to the gravity vector, with time on the radial axis and orientation angle on the circumferential axis; error bars, s.e.m.

To maintain constant temperature and minimise the pH gradient generated by electrolysis, the liquid medium was continuously circulated in a closed loop between the V-box, a cooled water bath and a 2-litre reservoir bottle (Supplemental Fig. S1 and Methods). In each V-box, the primary roots of five seedlings were imaged every 10 minutes with a camera mounted in front of the V-box (Supplemental Fig. S1 and Methods). We measured the actual field generated in the medium by immersing voltmeter probes in the two neighbouring positions of each seedling: as expected, the field measured in the medium was lower than the nominal imposed in air between the two electrodes, but still relatively uniform across the five positions (Supplemental Fig. S2a). At the same time, we measured the current passing through the circuit (Supplemental Fig. S2b and Methods).

To quantify the root response to the applied electric field, we took an image of the roots every 10 minutes and measured the angle between the root tip and the vertical gravity vector (Fig. 1b).

To confirm that the circulation of the liquid medium was effective in damping any pH gradient created by electrolysis and in eliminating any chemotropic effect, we imposed a field of 1.5 V/cm in a V-box without plants for one hour while maintaining liquid circulation. We then turned off the field and the liquid circulation, immediately positioned the plants in the V-box and imaged the roots for the next 80 minutes: we did not observe any significant deviation in root growth direction (t-test between 1.5 V/cm and 0 V/cm at 80 m, p=0.859), compared to roots in a V-box that never experienced the electric field (Supplemental Fig. S2c), indicating that no significant pH gradient was left in the medium.

### Root tip reorientation in external electric fields

We measured the response of wild-type (WT) *Arabidopsis* primary roots to a continuous electric field. The distribution of reorientations to a range of fields intensities between 0.5 V/cm and 2.5 V/cm shows a quick tropism towards the negative electrode, or cathode (Fig. 1c).

We wondered how much of this effect was due to trivial electrostatic, *i*.*e*. the physical pull towards the negative electrode due to a hypothetical net positive charge accumulated on the root tip, rather than a more complex biological response involving molecular signalling. To address this, we deactivated the roots by immersing them in a 50 °C bath for 10 min, until growth and gravitropism were suppressed (0/9 roots growing and bending to gravity after exposure to 50 °C, *vs*. 10/10 after exposure to 23 °C), transferred to the V-box and exposed them to a 2.0 V/cm electric field: the root response shows that this simple treatment was sufficient to completely inhibit electrotropism when compared to roots kept at a standard 23 °C temperature (t-test between 50 °C and 23 °C at 5 h, p<0.001), strongly suggesting that electrostatic alone could not explain the root response and that this is in fact a biological phenomenon (Supplemental Fig. S3).

### Response curves

The progressively sharper root tip reorientation as the field intensity was increased (Fig. 1c) suggests that the sensing mechanism is not acting as a simple on/off switch but that it can distinguish electric fields of different strengths. To quantitate this, we plotted the orientation angle at 5 h (“response”) as a function of the electric field intensity (“stimulus”), and found a best fit with a power function with exponent 0.45 (Fig. 2a, left), indicating that the resolution of the sensor is higher at low-intensity stimuli (steep response curve) than at high-intensity stimuli (shallow response curve). The analogous response curve as a function of the measured current intensity is best fit with a power function with exponent 0.33 (Fig. 2a, right).

**Figure 2.**
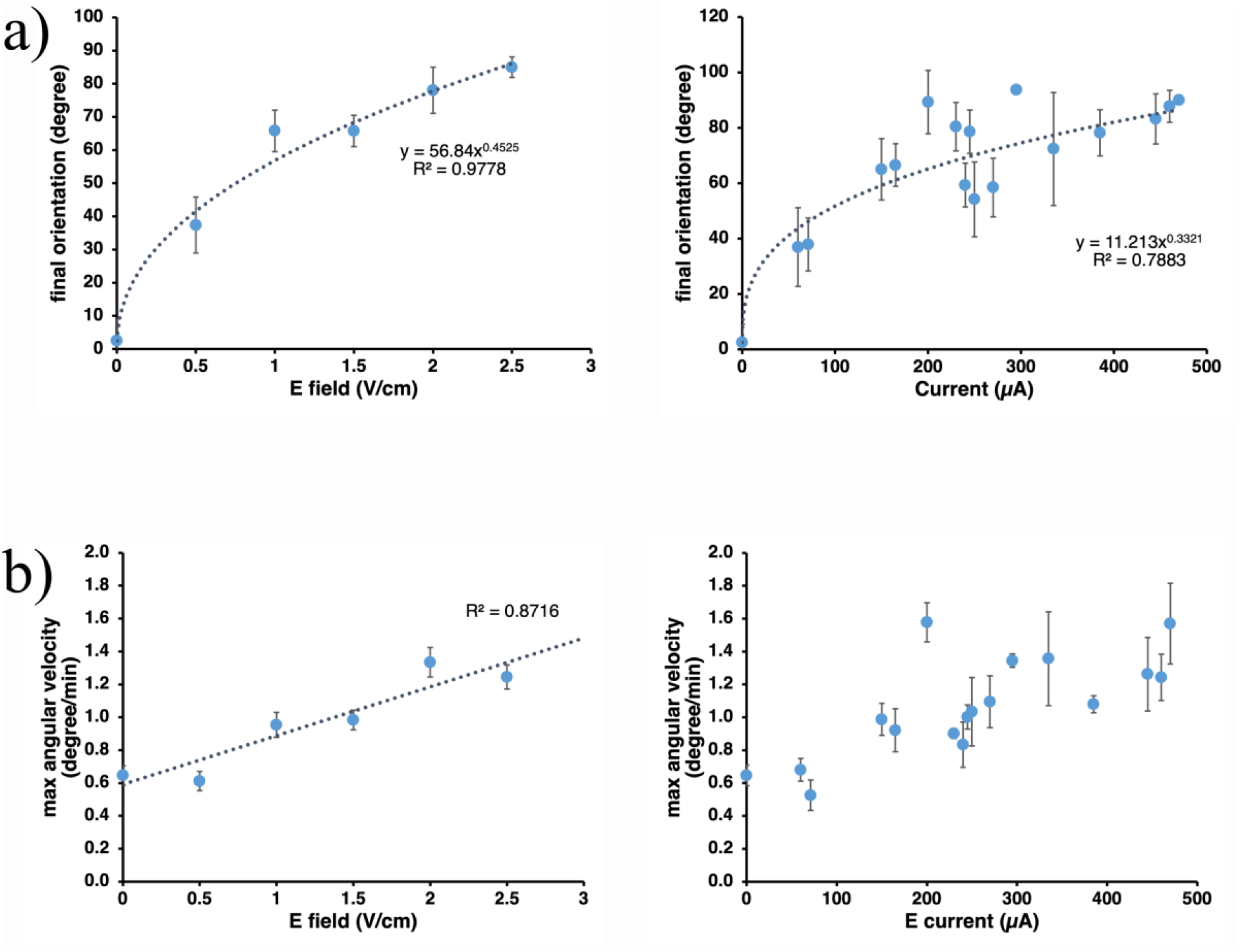
**(a)** Root electrotropism response curves: average root tip orientations after 5 h of exposure to a given electric field (left panel) or current (right panel); error bars, s.e.m.; R^2^, coefficient of determination. **(b)** Average maximum angular velocity reached by the root tip after 5 h of exposure to a given electric field (left) or current (right); error bars, s.e.m.; R^2^, coefficient of determination.

Since the root tip reorientations did not show any obvious overshoot (Fig. 1c), we looked more closely at the angular velocity: the maximum average velocity in WT roots was reached when the root tip was between 10 and 20 degrees orientation to the gravity vector and then progressively decreased as the root tip approached its maximum reorientation (Supplemental Fig. S4a). This indicates that the mechanism is able to sense and respond differently to the changing relative orientation of the tip with the external electric field, and to progressively slow down the root tip rotation as it approaches the target orientation. Moreover, the maximum angular velocity appears to be roughly proportional to the electric field strength (Fig. 2b, left), although this is much less evident as a function of the electric current (Fig. 2b, right).

### Root tips are not damaged

Early reports noted that protracted exposure to external fields could cause physical damage to plant root tips (Wawrecki and Zagórska-Marek, 2007). To control whether this was the case in our experimental conditions, we developed a simple chambered slide (V-slide) to be mounted on a standard confocal microscope stage (Supplemental Fig. S5a and Methods) with electrodes on the chamber’s sides and a circulating liquid medium for temperature control similar to the one implemented in the V-box (Supplemental Fig. S5b). Seedlings from the *Arabidopsis* transgenic line constitutively expressing the yellow fluorescent cell-membrane marker WAVE 131Y (Geldner et al., 2009) were mounted on the V-slide and imaged at cellular resolution while exposed to a 1.0 V/cm electric field (Methods). The field of view in our time-lapse images comprises the meristem, the transition zone and the distal elongation zone, but no cellular pattern perturbation was noticeable in any of these regions when compared to roots not exposed to the field (Supplemental Fig. S6). Also, the time-lapse suggests that asymmetric cell expansion in the elongation zone is causing the bending, as expected from other examples of root tropism (Gilroy, 2008).

To further confirm that roots exposed to the electric field are not damaged and maintain gravitropic response, we monitored root tips for 2 h after the electric field had been turned off, observing a clear gravitropic behaviour (Supplemental Fig. S7).

### Regions of competence

Since the root bending occurs in the elongation zone, we wanted to identify the region responsible for sensing the electric field. We excised distal fragments of the root at 125 μm, 300 μm, 400 μm and 500 μm from the tip (Fig. 3a), and then exposed the cut root to 1.5 V/cm (Methods).

**Figure 3.**
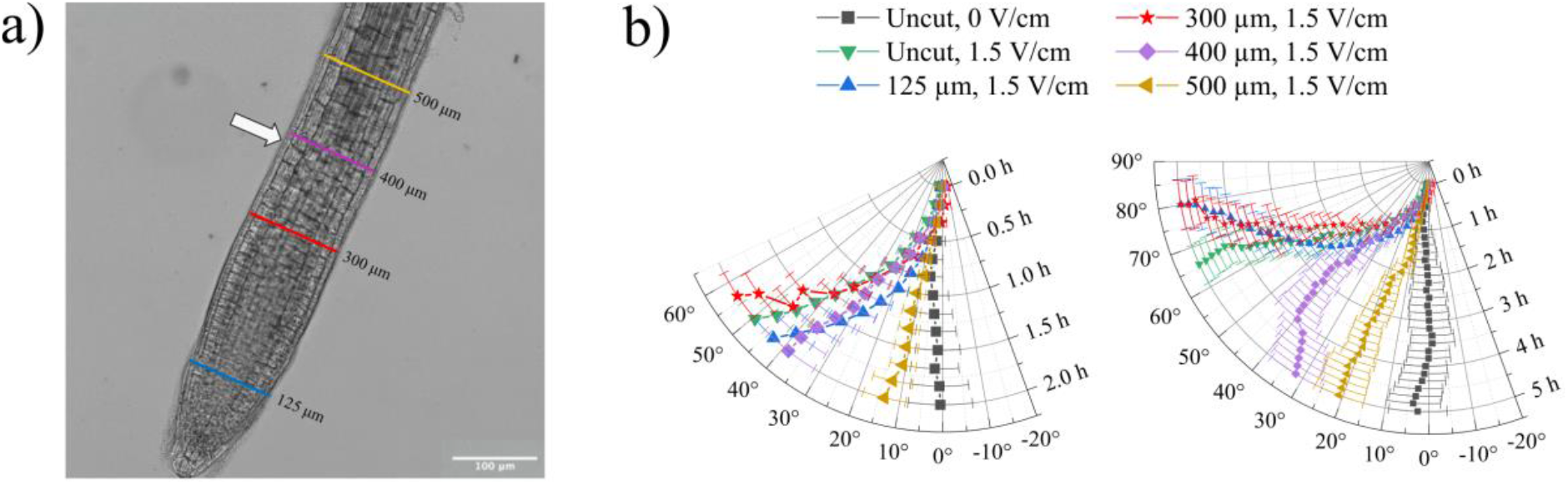
**(a)** Points of excision overlapped to a representative microscope image of the *Arabidopsis* primary root tip; arrow, transition point; scale bar, 100μm. (**b)** Polar plot of the average root tip orientations with respect to the gravity vector, with time on the radial axis and orientation angle on the circumferential axis; the same data is presented for the first 2 h (left panel) and for the full 5 h (right panel); error bars, s.e.m.

In the first 2 h (Fig. 3b left), roots cut at 400 μm from the tip turned to angles indistinguishable from uncut roots exposed to the same field (Wilcoxon between 400 μm cut and uncut at 1.5 V/cm at 2 h, p=0.211), while roots cut at 500 μm from the tip did not turn and at 2 h showed orientations indistinguishable from uncut roots not exposed to a field (t-test between 500 μm cut at 1.5 V/cm and uncut at 0 V/cm at 2 h, p=0.164). These results indicate that the 400 μm distal fragment is not necessary for the early (2 h) electrotropic response, while the 500 μm distal fragment is; since the elongation zone involved in the bending spans an extended region proximal to the 500 μm cut, we conclude that in *Arabidopsis* roots the region between 400 μm and 500 μm from the tip is necessary for early sensing an external electric field.

Between 2 h and 5 h of exposure (Fig. 3b right), while roots cut at 300μm on average continue to turn like the uncut roots (t-test between 300 μm cut and uncut at 1.5 V/cm at 5 hrs, p=0.071), roots cut at 400 μm quickly fail to sustain their response and at 5 h they show tip orientations on average different than those of uncut roots exposed to the same field (t-test between 400 μm cut and uncut at 1.5V/cm at 5 hrs, p<0.001). These results indicate that the 300 μm distal fragment is not necessary to maintain the electrotropic response up to 5hrs, while the 400 μm distal fragment is; we conclude that in *Arabidopsis* roots the region between 300 μm and 400 μm from the tip is necessary for prolonged sensing of the imposed electric field.

Interestingly, previous studies suggested that the movement of Ca++ ions accumulated in the mucilage at the very tip of the root might be involved in the electrotropic sensing (Marcum and Moore, 1990). Since any excision above 125 μm from the tip essentially removes all the mucilage from the root, our results disprove this hypothesis for *Arabidopsis*.

### Auxin distribution is not altered by the electric field

The fact that roots without tips could still respond to the electric field shows that even a major disruption of the stereotypical auxin redistribution mechanism is not sufficient to inhibit electrotropism.

On the other hand, an asymmetric accumulation of auxin is required for asymmetric cell elongation and root bending in some tropisms, as suggested by the classic Cholodny-Went model (Thimann and Went, 1937). To test whether this model applied to electrotropism, and whether the external electric field is sufficient to induce an asymmetric distribution of auxin in the root, we exposed roots expressing the auxin-sensitive fluorescent reporter *R2D2* (Liao et al., 2015) to a field of 1.0 V/cm for 30 minutes, before quickly mounting them on a microscope slide and imaging them with a confocal microscope (Methods). A ratiometric quantification of *R2D2* signal (Kral et al., 2016) in each epidermal cell (Fig. 4a and Methods) showed that the average auxin response measured in the epidermal cells on the side facing the negative electrode and on the side facing the positive electrode were statistically indistinguishable (Fig. 4b; Methods), both in the distal (Wilcoxon test, p=0.696) and the proximal (Wilcoxon test, p=0.843) region of the root, indicating that auxin is not asymmetrically distributed after exposure to the electric field.

**Figure 4.**
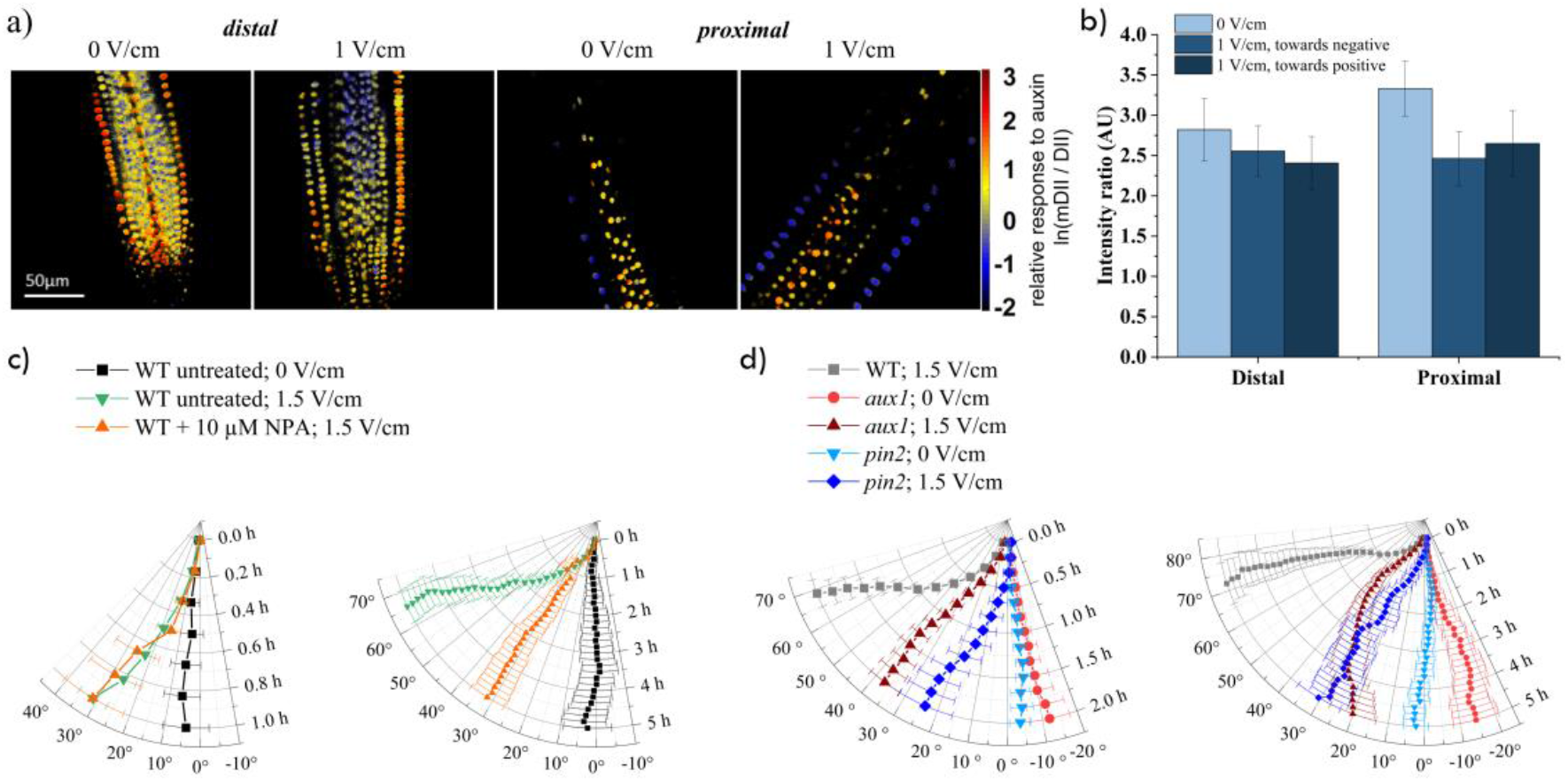
**(a)** Relative response to auxin concentration in *Arabidopsis* primary root tips, indicated by the ratiometric fluorescent reporter R2D2 (see Methods); left panels: representative tip (distal) regions at 0 V/cm and 1 V/cm; right panels: representative higher (proximal) regions at 0 V/cm and 1 V/cm; scale bar, 50μm. (**b)** Quantification of the average relative response to auxin (as shown in a) in the epidermis cell layer facing the positive or negative electrode, in the distal and proximal regions; the data for 0 V/cm is an average between the two sides; error bars, s.e.m. **(c-d)**, Polar plots of the average root tip orientation relative to the gravity vector, with time on the radial axis and orientation angle on the circumferential axis: NPA treatment **(c)**; *aux1* and *pin2* mutants **(d)**; error bars, s.e.m.

Moreover, both sides were also indistinguishable from the average between the two sides in roots not exposed to the field (mock), both in the distal (Wilcoxon test between the negative exposed side and mock, p=0.775; and between the positive exposed side and mock, p=0.881) and in the proximal (Wilcoxon test between the negative exposed side and mock, p=0.041; and between the positive exposed side and mock, p=0.098) regions.

### Auxin transport is not necessary for early response

Previous work indicated that auxin transport inhibitors can inhibit electrotropism response in maize roots (Ishikawa and Evans, 1990). To verify this in *Arabidopsis*, and further explore the role of auxin in root electrotropism, we used N-1-naphthylphthalamic acid (NPA) to inhibit polar auxin transport (Sabatini et al., 1999) while exposing the roots to the electric field. We tested 0.1 μM, 1.0 μM and 10 μM NPA and found that 10 μM NPA was the lowest concentration of NPA to inhibit gravitropism (Methods), a known auxin-dependent tropism.

We pre-treated the seedlings for 3 hours in liquid medium containing 10 μM NPA (Methods) and then transferred them to a V-box containing 10 μM NPA in the medium and the reservoir. In the first 1 hour of field exposure (Fig. 4c left) the NPA-treated roots reoriented at angles indistinguishable from those of untreated ones (Wilcoxon test between NPA-treated and untreated at 1 h, p=0.350). From 1 to 5 hours of exposure (Fig. 4c right), NPA-treated roots respond less than the untreated (t-test between NPA-treated and untreated at 5 h, p<0.001), but still significantly more than the untreated and not exposed to the field (t-test between untreated at 1.5 V/cm and 0 V/cm at 5 h, p=0.001). These results indicate that auxin polar transport is not necessary for an early electrotropic response, but might play a role in maintaining a long-term orientation.

To further explore the role of auxin transport, we tested the mutants of PIN2, an auxin cellular exporter (Chen et al., 1998), and mutants of AUX1, an auxin cellular importer (Marchant et al., 1999). Since *pin2* and *aux1* mutants do not respond to gravity, to obtain a sufficient number of roots growing vertically in preparation for the electrotropic assay, we wrapped the sides and bottom of the nursery boxes with aluminium foil to induce negative root phototropism towards the bottom of the box (Methods). The same setup was used to germinate WT plants to be compared with these to mutants.

Roots of *pin2* mutants (Fig. 4d) showed a significant response (paired Wilcoxon between the time-points 0 h and 2 h, p<0.01; and between the time-points 0 h and 5 h, p<0.01), but weaker than WT (t-test between WT and *pin2* at 2 h, p<0.01; and between WT and *pin2* at 5 h, p<0.01) in the same conditions. Their angular velocity did not show an obvious decrease before reaching the target orientation, as with WT roots (Supplemental Fig. S4b).

Analogously, roots of *aux1* mutants (Fig. 4d) showed a significant early response (paired t-test between the time-points 0 h and 2 h, p<0.01), but weaker than WT (t-test between WT and *pin2* at 2 h, p<0.01; t-test between WT and *pin2* at 5 h, p<0.01) in the same conditions. Interestingly, *aux1* roots on average failed to maintain their orientation for longer period (paired t-test between the time-points 0 h and 5 h, p=0.051). Moreover, their angular velocity on averaged decreased while approaching the final orientation, as with WT roots (Supplemental Fig. S4b).

Taken together, these results suggest that although auxin transport seems to play a role in maintaining a sustained response to the electric field, it is not necessary in triggering early electrotropism.

### Cytokinin biosynthesis is necessary for electrotropism

Given our results suggesting a limited role for auxin during electrotropism, we wondered which other plant hormones might be involved instead. Root hydrotropism, or the growth towards high concentration of water, was also previously shown to be largely independent from auxin distribution in *Arabidopsis* (Shkolnik et al., 2016), while it requires biosynthesis and asymmetric distribution of cytokinin (Chang et al., 2019). Drawing an analogy with hydrotropism, we considered the possibility that electrotropism might act through cytokinin as well.

To test this hypothesis, we analysed root electrotropism in triple mutants of *AtIPTs*, a family of adenosine phosphate-isopentenyltransferases required for the first step of isoprenoid cytokinin biosynthesis (Miyawaki et al., 2006; Kamada-Nobusada and Sakakibara, 2009). Within the tested mutants, although a high degree of redundancy is expected among the *AtIPTs*, we found two distinct behaviours: the triple mutants *atipt1,3,5* and *atipt3,5,7* both responded strongly to the electric field (Fig. 5a) (t-test between 1.5 V/cm and 0 V/cm at 5 h, p<0.001 in both cases), while the triple mutants *atipt1,3,7* and *atipt1,5,7* responded well in the first 2 h of exposure (Fig. 5b, left) (t-test between 1.5 V/cm and 0 V/cm at 2 h, p<0.001 for both mutants) but showed a much weaker, although still significant, response at 5 h (Fig. 5b right) (t-test between 1.5 V/cm and 0 V/cm at 5 h, p<0.01 for *atipt1,5,7* and p<0.001 for *atipt1,3,7*; t-test between WT and mutant both at 1.5 V/cm at 5 h, p<0.001 for *atipt1,5,7* and p<0.01 for *atipt1,3,7*). These results suggest that cytokinin biosynthesis is in part required for long-term root electrotropism, and that for this phenotype *AtIPT1 and AtIPT7* dominate their family in a redundant way (the response is the weakest when both are mutated).

**Figure 5.**
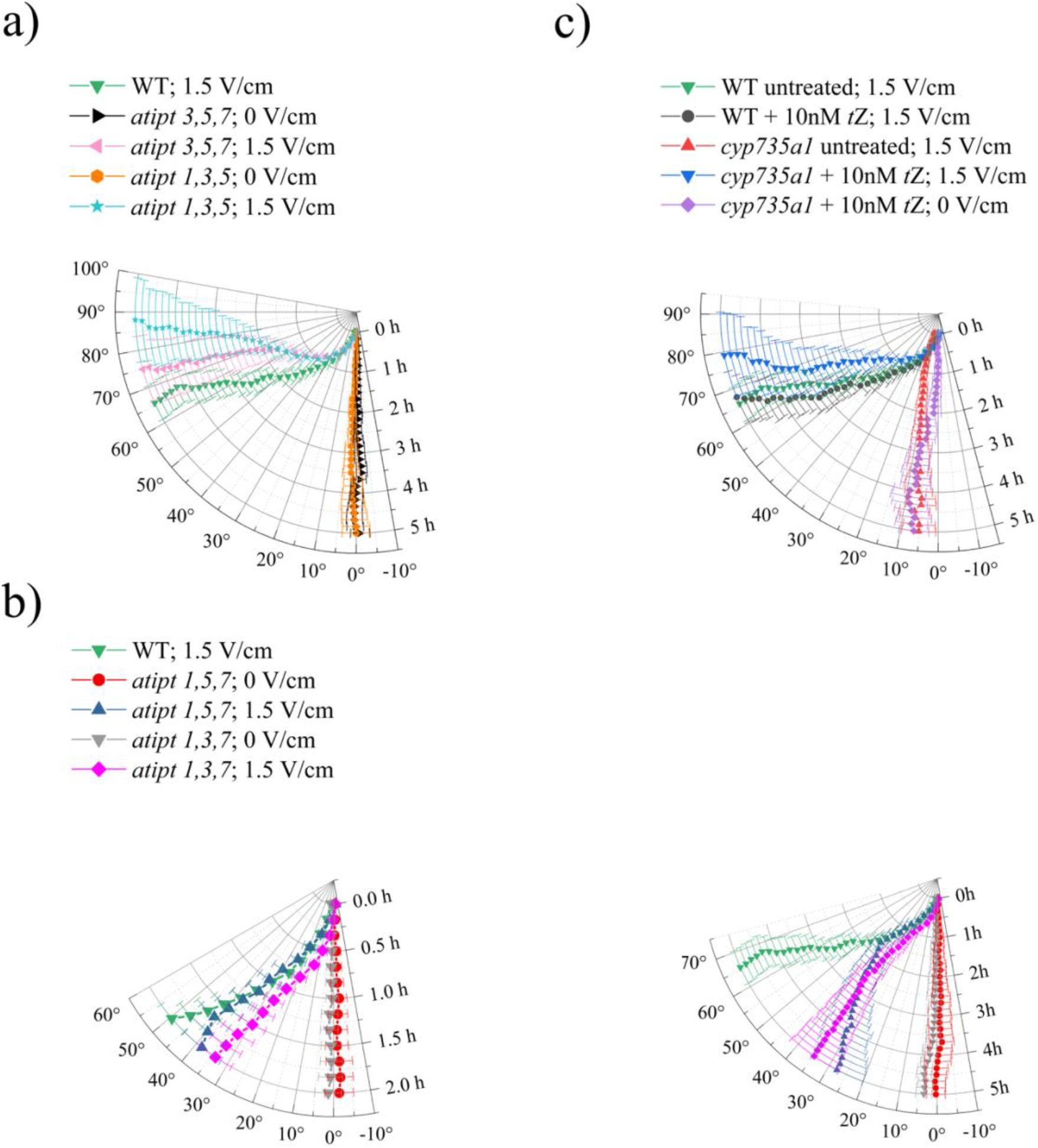
Polar plots of the average root tip orientation relative to the gravity vector, with time on the radial axis and orientation angle on the circumferential axis: **(a)** *atipt* triple mutants with strong electrotropic response; **(b)** *atipt* triple mutants with weak electrotropic response; **(c)** *cyp735a1* mutant; error bars, s.e.m.

To confirm the role of cytokinin, we also tested the requirement for *CYP735A1*, a cytochrome P450 monooxygenase enzyme acting downstream of *AtIPTs* and necessary for the biosynthesis of the *trans*-zeatin (*t*Z) variation of cytokinin (Takei et al., 2004). Interestingly, roots of *cyp735a1* mutants completely failed to respond to the electric field (Fig. 5c), with tip orientations indistinguishable from that of WT roots not exposed to the field (t-test between *cyp735a1* exposed to 1.5 V/cm and WT not exposed at 5 h, p=0.748). Crucially, this phenotype could be rescued with the addition of the cytokinin *trans-*Zeatin (*t*Z) to the medium (Fig. 5c) (t-test between *cyp735a1* +10nM *t*Z at 1.5 V/cm and 0 V/cm, p<0.001), while the same treatment did not affect WT response (Fig. 5c).

Taken together, these results indicate that cytokinin is necessary for electrotropism in *Arabidopsis* roots.

To further investigate a possible parallel between the molecular pathways involved in electrotropism and hydrotropism, we tested mutants of *MIZU-KUSSEIN1 (MIZ1)*, which is necessary for hydrotropism (Kobayashi et al., 2007) and its functional cytokinin asymmetric distribution in roots (Chang et al., 2019). Translational fusion reporters have shown MIZ1 localisation in the lateral root cap and the cortex of meristem and elongation zone (Dietrich et al., 2017): although the root tip is not necessary for hydrotropic response (Dietrich et al., 2017), this requires MIZ1 in the transition zone (Dietrich et al., 2017).

When we exposed roots of *miz1* mutants to 1.5 V/cm (Fig. 6) they showed an unperturbed electrotropic response in the first 2 h (Wilcoxon test between *miz1* and WT at 2 h, p=0.859) and perhaps a weakened response at 5 h, although with only a weak statistical significance (t-test between *miz1* and WT at 5 h, p=0.033), indicating that MIZ1 is not necessary for early root electrotropism.

**Figure 6.**
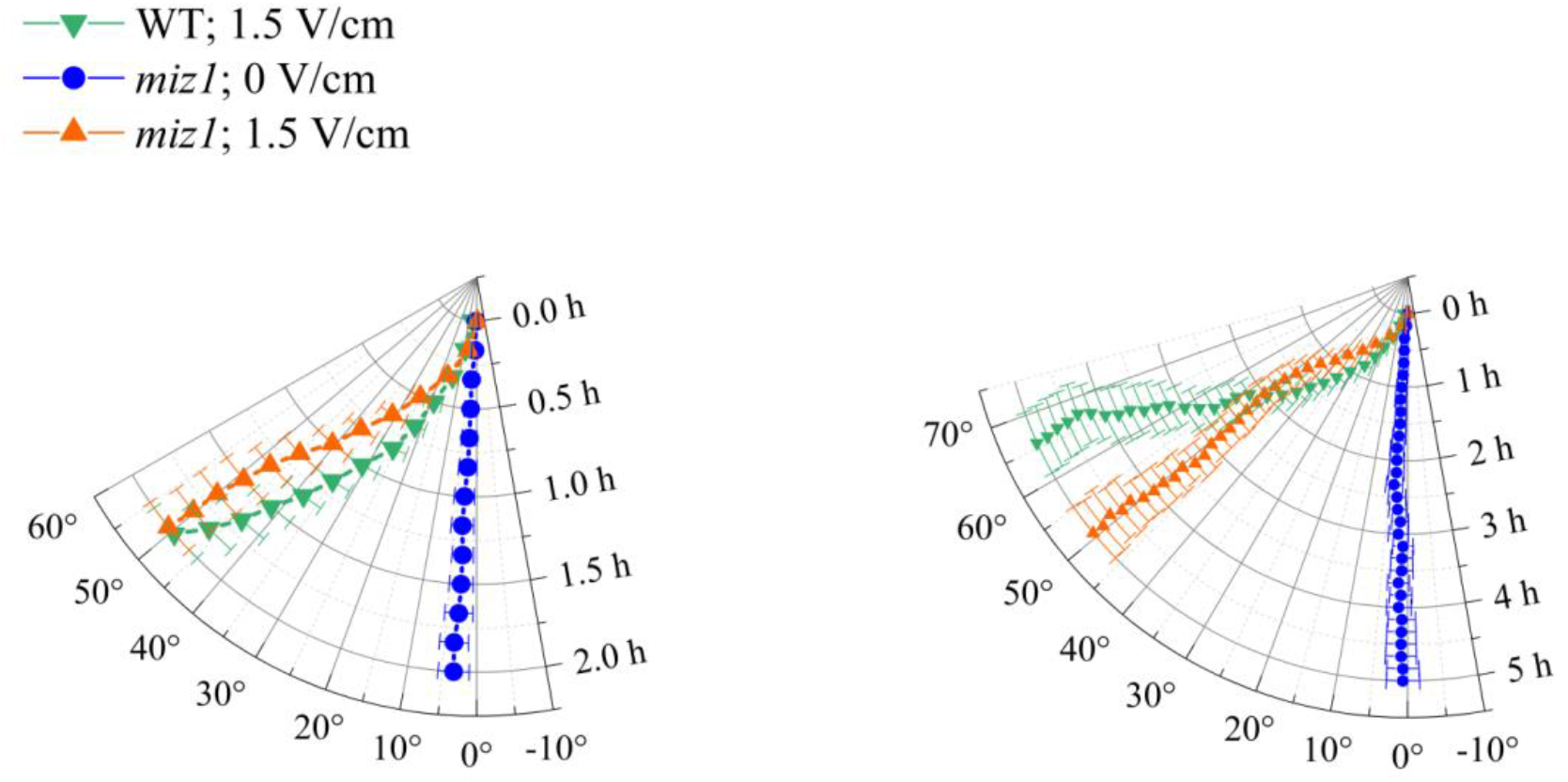
Polar plots of the average root tip orientation in *miz1* mutant relative to the gravity vector, with time on the radial axis and orientation angle on the circumferential axis.

Taken together, these surprising results indicate that cytokinin plays an important role in root electrotropism, but that the underlying molecular pathway differs early on from that of hydrotropism.

## DISCUSSION

The capability of plant roots to sense and combine numerous physical and chemical signals in soil is quite extraordinary, notably in the absence of a centralised processing system. The more we understand about the physical and molecular mechanism behind the various root tropisms, the closer we will get to a complete understanding of signal integration in plants. In this work we focused on the little-studied phenomenon of root electrotropism and present its first quantitative characterisation in the flowering plant model system *Arabidopsis thaliana*.

A non-trivial observation from this work is that plant roots respond to weak (order of 1 V/cm and 100 μA) external electric fields and currents in a progressive way, with an increasingly stronger tendency to align with the field as the field and current intensities increase. This is interesting, because it reveals a sophisticated way to discriminate between highly charged particles (*e*.*g*. ions, micro-organisms, other plant roots) and weakly charged ones. At the same time, the absence of overshooting also points to a mechanism that perhaps can be modelled along the lines of a proportional–integral–derivative control system (Chevalier et al., 2019). Moreover, we show that the kinetics (response curve) of root electrotropism follows a power-law with exponents similar to those that have been traditionally measured in physiological responses to external stimuli in animal systems (Stevens, 1970): our results suggest that a power-law response to external stimuli might be a universal feature across life kingdoms, and that it does not require a nervous system. Moreover, it might reveal constraints on the genetic architecture of the underlying sensory system (Adler et al., 2014).

A second, perhaps unexpected, result from this work is the identification of two regions in *Arabidopsis* roots required for the detection of the field: the section between roughly 400 μm and 500 μm from the tip is necessary for the early detection (within 2 h from exposure), while the section between roughly 300 μm and 400 μm is necessary for a prolonged detection (up to 5 h from exposure). Interestingly, these sections seem to correspond to the well-characterised anatomical transition zone, between the meristem and the elongation zone in roots, which has been previously involved in root sensing (Muthert et al., 2020). In future works, it will be crucial to discern whether the electrotropism mechanism depends on a relatively large region like the transition zone or if it can be narrowed to a more specific cell population therein.

Finally, we present evidence against the assumption that auxin asymmetric distribution is required for electrotropism in *Arabidopsis*, similarly to what found for hydrotropism (Shkolnik et al., 2016) and phototropism (Kimura et al., 2018). Instead, we show that cytokinin is required for a full electrotypic response, although its response-specific regulation seems to act through a different pathway than the one established for hydrotropism.

Overall, our results show that root electrotropism requires a sensing mechanism likely localised in the transition zone, a limited role for auxin but an important role for cytokinin. The latter could be involved in a signalling or regulatory mechanism in the transition zone, in an actuator mechanism (tissue bending) in the elongation zone, or both.

Cytokinins have been shown to regulate root meristem activity and size by controlling cell proliferation (Beemster and Baskin, 2000) and the developmental progression from proliferation to elongation in the transition zone (Ioio et al., 2007). It has been suggested that root bending in hydrotropism is based on asymmetric distribution of cytokinin in the meristem to induce asymmetric cell proliferation (Chang et al., 2019). This seems an unlikely mechanism for electrotropism, because roots without meristem (a 400μm segment cut from the tip) still respond to the electric field, suggesting that root bending in this case does not depend much on cell proliferation. An alternative mode of action for cytokinins during electrotropism could be based on its regulatory action in the transition zone, where an asymmetric delay in elongation would result in root bending. Although we have shown that MIZ1 is not required for root electrotropism, it is still possible that a MIZ1-independent mechanism could generate an asymmetry in cytokinin distribution in the transition zone.

More recently, cytokinins have also been involved in stress response through regulation of downstream factors and through crosstalk with other hormones (Li et al., 2021; Wu et al., 2021). This signalling role could be relevant during elecrotropism, especially since the transition zone has often been associated with signal integration in the root (Baluška et al., 2010).

Future work on root electrotropism should focus on testing these hypotheses regarding the role of cytokinin and on illuminating the still unknown molecular mechanism involved in sensing an electric field or current.

## METHODS

### Plant material

Wild-type and mutant *Arabidopsis thaliana* plants were all from the Columbia (Col-0) ecotype; the following mutant alleles were used: *aux1-7* (NASC id 9583) for *aux1*; *eir1-1* (NASC id 8058) for *pin2;* SALK_093028C (NASC id N654306) for *cyp735a1, miz1-1* for *miz1* (courtesy of Prof. A. Kobayashi); *atipt* triple mutants as previously described with *atipt1-1, atipt3-2, atipt5-2, atipt7-1* (Miyawaki et al., 2006) (courtesy of Prof. O. Leyser).

The fluorescent line *WAVE131Y* is expressing *pUBQ10::WAVE131:YFP* (NASC id N781535); the fluorescent line *R2D2* is expressing *RPS5A-mDII-ntdTomato, RPS5A-DII-n3xVenus* (courtesy of Dr. Teva Vernoux).

Seeds were imbibed in water and kept in the dark for 2 days at 4 °C, in order to synchronise germination. All seeds were surface sterilised using 50% Haychlor bleach and 0.0005% Triton X-100 for 3 minutes and then rinsed 6 times with sterilised milliQ water. Seed germination protocols are described in the experiment-specific Methods sections below. Unless otherwise specified, all experiments were conducted with primary roots of seedlings 5-8 days post-germination, with roots approximately of the same length to be in the field of view of the V-box camera.

### Electrotropism assay (V-box)

Seeds were sown individually inside PCR tubes filled with 1X MS gel medium: 0.44% Murashige and Skoog (MS) Basal medium (Sigma-Aldrich, M5519), 0.5% sucrose, 0.05% MES hydrate (Sigma-Aldrich M8250), 0.8% agar (Sigma-Aldrich 05040), pH adjusted to 5.7 with TRIS HCl (Fisher-Scientific 10205100). The PCR tubes had their end cut out to allow the root to grow through, and placed in a 3D-printed (Ultimaker 2+) holder inserted in a Magenta box (Sigma-Aldrich V8380). The Magenta box was filled with 150 ml of 1/500X MS liquid medium (0.00088% MS Basal medium, 0.5% sucrose, 0.05% MES hydrate, pH adjusted to 5.7 with TRIS HCl) to reach the end of the PCR tubes. These germination, or “nursery”, boxes were placed in a growth chamber at 22 °C, with a 16 h/8 h light/dark photoperiod and light intensity 120 μmol m^-2^ s^-1^

In preparation for the electrotropism assay, each PCR tube containing a single seedling was transferred to a modified 3D-printed holder in a Magenta box filled with 1/500X MS liquid medium. The modified module consisted of a main body with five holes for the PCR tubes containing the seedlings, and two side clips to position the electrodes consisting of platinum-iridium (Platinum:Iridium = 80:20; Alfa Aesar 41805.FF) foils (Fig. 1a), which were connected to an external power supply. In this paper, we refer to the Magenta box and its holder as the “V-box”. In addition to the five holes for the plants, four extra holes were designed: two at each end of the front side to pump the medium out of the V-box right on top of the two electrodes, and two on the back to pump the medium in, using a tubing system and peristaltic pumps (Verdeflex AU R2550030 RS1) to circulate the medium at a speed of 1 ml/sec to and from a 2 l reservoir bottle (Extended Data Fig. 1). This configuration was designed and tested to ensure a slow and symmetric flow of medium in the box, eliminating any biased effect of the flow on the roots, which was confirmed by the control experiments at 0 V/cm, performed with the pump circulating the medium at the same speed as in the exposure experiments. Crucially, between the V-box and the reservoir bottle, the tubings were immersed in a cooled water bath (Grant Instruments, LTDGG) maintained at the constant temperature of 19 °C (Extended Data Fig. 1), which was enough to maintaining the medium inside the V-box at a constant 22 °C, as measured. All electrotropic experiments were performed at constant illumination.

The electric field was generated with a power supply attached to the platinum-iridium foil electrodes that were immersed in the liquid medium in the V-box. Standard electric wires were soldered on the top of two electrodes, always kept outside the liquid medium to avoid contaminants from the solder. The voltage was set constant on the power supply, while the current was measured independently with a multimeter in series.

In order to record the movement of the roots over time, a Raspberry Pi camera module V2 (913-2664) connected to a Raspberry Pi board module B+ (137-3331) was used. The Raspberry Pi was programmed to take a picture every 10 minutes, using the command *crontab* in the local Raspbian OS.

WT electrotropism assays (Fig. 1) were performed with the following sample sizes N and number of replicates R: 0 V/cm, N=10, R=2; 0.5 V/cm, N=9, R=2; 1.0 V/cm, N=8, R=2; 1.5 V/cm, N=21, R=5; 2.0 V/cm, N=18, R=6; 2.5 V/cm, N=20, R=7; foil-wrapped and 1.5 V/cm (control for *pin2* and *aux1*), N=15, R=3.

Mutants electrotropism assays (Fig. 4,5,6) were performed with the following sample sizes N and number of replicates R: *aux1* at 1.5 V/cm, N=18, R=4; *aux1* at 0 V/cm, N=14, R=3; *pin2* at 1.5 V/cm,, N=13, R=4; *pin2* at 0 V/cm,, N=13, R=3; *atipt3,5,7* at 1.5 V/cm, N=15, R=3; *atipt3,5,7* at 0 V/cm, N=10, R=2; *atipt1,3,5* at 1.5 V/cm, N=15, R=3; *atipt1,3,5* at 0 V/cm, N=10, R=2; *atipt1,5,7* at 1.5 V/cm, N=14, R=3; *atipt1,5,7* at 0 V/cm, N=10, R=2; *atipt1,3,7* at 1.5 V/cm, N=14, R=5; *atipt1,3,7* at 0 V/cm, N=10, R=2; *cyp735a1* at 1.5 V/cm, N=20, R=4; *miz1* at 1.5 V/cm, N=9, R=2; *miz1* at 0 V/cm, N=10, R=2

The control for medium circulation efficiency (Fig. S2) was performed with the following sample sizes N and number of replicates R: 1.5 V/cm, N=10, R=2; 0 V/cm, N=10, R=2.

The experiment showing gravitropism after 2 hours of electric field (Fig. S7) was performed with a sample size N=10 and replicates R=2.

### High temperature treatment

Wild-type Col-0 seeds were germinated and grown in the nursery boxes as described in the Electrotropism Assay section. The boxes undergoing treatment were then immersed in a water bath set at 50 °C, for 10 minutes. The PCR tubes containing the seedlings were then transferred to a V-box, exposed to an electric field of 2.0 V/cm for 5 hours and imaged, as described in the Electrotropism Assay section.

Electrotropism assays for treated roots were performed with the following sample sizes N and number of replicates R: 23°C, N=18, R=6; 50°C, N=10, R=2.

For the gravitropism control, seedlings were treated with high temperature as described above and then transferred on 1X MS agar plates, to complete the assay as described in the Gravitropic Assay section.

Gravitropic assays for treated roots were performed with the following sample sizes N and number of replicates R: 23°C, N=10, R=2; 50°C, N=9, R=1).

### Electrotropism on microscope (V-slide)

Seeds of the transgenic reporter line *WAVE131Y* (see Plant Material section) were germinated on 1X MS agar medium (0.44% MS Basal medium, 0.5% sucrose, 0.05% MES hydrate, pH adjusted to 5.7 with TRIS HCl, 0.8% agar) in square plates kept vertical in a growth chamber at 22 °C, with a 16 h/8 h light/dark photoperiod.

At 2 days post-germination the seedlings were mounted on the V-slide with 1/500X MS liquid medium (0.00088% MS Basal medium, 0.5% sucrose, 0.05% MES hydrate, pH adjusted to 5.7 with TRIS HCl). The V-slide was then connected to the medium perfusion system circulating the same 1/500X MS medium, and the V-slide’s electrodes were connected to the power supply (Voltcraft PS-1302-D).

Imaging was performed on a Leica SP5 laser scanning confocal microscope with 20X air objective, while the liquid medium was circulated and the electric field maintained constant. Fluorophore was excited with the 514 nm Argon laser line and the emission collected with PMT detector at 524-570 nm. Images were collected at intervals of 2 minutes.

### Root tip excisions

Seeds were germinated in nursery boxes as described.

3 days post-germination seedlings still in their PCR tubes, as previously described, were transferred to hard 1X MS agar medium (0.44% MS Basal medium, 0.5% sucrose, 0.05% MES hydrate, pH adjusted to 5.7 with TRIS HCl, 5.0% agar) where the roots were manually dissected using a dental needle (Sterican, 27G) under a dissecting microscope (Nikon SMZ1000) at 180X magnification, following the published protocol (Kral et al., 2016).

After root tip excision, the PCR tubes containing the seedlings were immediately moved into the V-box for the electrotropism assay.

Electrotropism assays on excised roots were performed with the following sample sizes N and number of replicates R: 125 μm, N=16, R=4; 300 μm, N=7, R=2; 400 μm, N=10, R=2; 500 μm, N=10, R=2; uncut at 0 V/cm, N=10, R=2; uncut at 1.5 V/cm, N=21, R=5.

### R2D2 reporter

Seeds of the transgenic reporter line *R2D2* (see Plant Material section) were germinated in nursery boxes and at 3days post-germination exposed to 1.0 V/cm for 30min in V-boxes, as described in the Electrotropism Assay section. After exposure, *R2D2* roots were quickly mounted on standard microscope slides with sterile deionised water and imaged using Leica SP5 laser scanning confocal microscope, with 63X water immersion objective.

*R2D2* expresses two versions of a protein that forms a complex with auxin: DII-n3xVENUS, which is degraded within minutes upon binding with auxin; mDII-ntdTOMATO, which contains a modified, non-degradable, version of DII. Both of the proteins are localised in the nucleus. We followed the published protocol (Liao et al., 2015) to separately collect the emission from mDII-ntdTOMATO (Extended Data Fig. 6a) and DII-n3xVENUS (Extended Data Fig. 6b).

For each root, a mean background was defined as the average pixel intensity in the mDII channel of a 80×80 pixels corner of the field of view not occupied by the root. The mean background was then subtracted form all pixel intensities in both channels (mDII and DII). We manually segmented with FIJI (Schindelin et al., 2012) the most visible cell nuclei in the epidermis of each root, both in the distal (12 roots in mock conditions and 16 roots exposed to the field) and in the proximal (11 roots in mock conditions and 13 roots exposed to the field) regions of the root tip, and quantified the average pixel intensity (corrected after background subtraction) for each segmented nucleus. Finally, we calculated the natural log of the ratio between the average mDII (non-degradable, auxin-independent) and DII (degradable, auxin-dependent) signals in each nucleus and mapped it on top of the root image (Extended Data Fig. 6d).

For each root, we calculated the average and standard deviation of these ratios, for epidermal nuclei facing the anode or the cathode (Fig. 4b).

Sample size as following: exposed to E field and imaged in distal region, N=16; exposed to E field and imaged in the proximal region, N=13; not exposed to E field and imaged in the distal region, N=24; not exposed to E field and imaged in the proximal region, N=22.

### NPA treatment

To find the minimum concentration of N-1-naphthylphthalamic acid (NPA) that inhibits gravitropism, seeds were germinated on 1X MS agar medium (0.44% MS Basal medium, 0.5% sucrose, 0.05% MES hydrate, pH adjusted to 5.7 with TRIS HCl, 0.8% agar) and at 5-8 days post-germination were transferred for 3 hrs into cell culture dishes containing 5 ml of 1/500X MS liquid medium (0.00088% MS Basal medium, 0.5% sucrose, 0.05% MES hydrate, pH adjusted to 5.7 with TRIS HCl) plus NPA at 0.1 μM, 1.0 μM and 10 μM.

To quantify the effect of NPA on electrotropism, after being treated with a concentration of 10 μM NPA for 3 hours as described above, the seedlings were transferred inside a V-box and exposed to a 1.5 V/cm electric field as described in the Electrotropism Assay section. Both the V-box and the reservoir bottle contained 10 μM NPA throughout the experiment.

Sample sizes N and number of replicates R were the following: untreated, N=21, R=5; 10 μM NPA treated, N=21, R=5.

### Cytokinin treatment

Both the 1X MS agar medium (0.44% MS Basal medium, 0.5% sucrose, 0.05% MES hydrate, pH adjusted to 5.7 with TRIS HCl, 0.8% agar) contained in the PCR tubes, and the 1/500X MS liquid medium (0.00088% MS Basal medium, 0.5% sucrose, 0.05% MES hydrate, pH adjusted to 5.7 with TRIS HCl) in the nurseries, were supplemented with 10nM *trans*-zeatin (Sigma-Aldrich Z0876) as previously suggested (Miyawaki et al., 2006). Still in their PCR tubes, seedlings were transferred into V-boxes, filled with 1/500X MS liquid medium supplied with 10nM *trans*-zeatin to conduct the electrotropism experiments.

Sample sizes N and number of replicates R were the following: WT + tZ, N=14, R=3; *cyp735a1* +tZ at 1.5 V/cm, N=13, R=3; *cyp735a1* +tZ at 0 V/cm, N=14, R=3.

### Gravitropism assay

In the high-temperature experiment, after the treatment the seedlings were transferred onto 1X MS square agar plates (0.44% MS Basal medium, 0.5% sucrose, 0.05% MES hydrate, pH adjusted to 5.7 with TRIS HCl, 0.8% agar).

In the NPA experiment, after the treatment the seedlings were transferred onto 1/500X MS square agar plates (0.00088% MS Basal medium, 0.5% sucrose, 0.05% MES hydrate, pH adjusted to 5.7 with TRIS HCl, 0.8% agar) containing the desired concentration of NPA.

In both cases, the plates were moved to a growth room (22 °C, 16 h/8 h light/dark photoperiod), rotated by 90 degrees to position the roots in a horizontal orientation, and monitored for root gravitropic response.

### Statistical analysis

When comparing two samples of measurements, each distribution was first checked for normality with the Shapiro-Wilk test with alpha-level = 0.05. Normal distributions were tested with the two-tails Student’s t-test without assuming equal variances (Welch two sample t-test); if one of the two distributions was not normal, the non-parametric Mann-Whitney U (Wilcoxon) test was used. Unless stated otherwise, all comparisons were performed assuming independence (unpaired test). All statistical tests were performed with R.

## ACKNOWLEDGEMENTS

We thank T. Vernoux for providing *R2D2* seeds; O. Leyser for providing *atipt* triple mutant seeds; A. Kobayashi and Tohoku University for providing *miz1-1* seeds; A. Guyon, O. Kranse, C. Keyzor, C.D. Livesey-Clare for generous technical help; M. Levin and I. Bordeu for critical discussions at various stages of the project. This work was partly supported by the BBSRC Impact Acceleration Account BB/S506667/1 and the Allen Discovery Center at Tufts University; N.K. was supported by the BBSRC DTP grant BB/M011178/1.

