## Supplemental Figures for "Root electrotropism in Arabidopsis does not depend on auxin distribution but requires cytokinin biosynthesis"

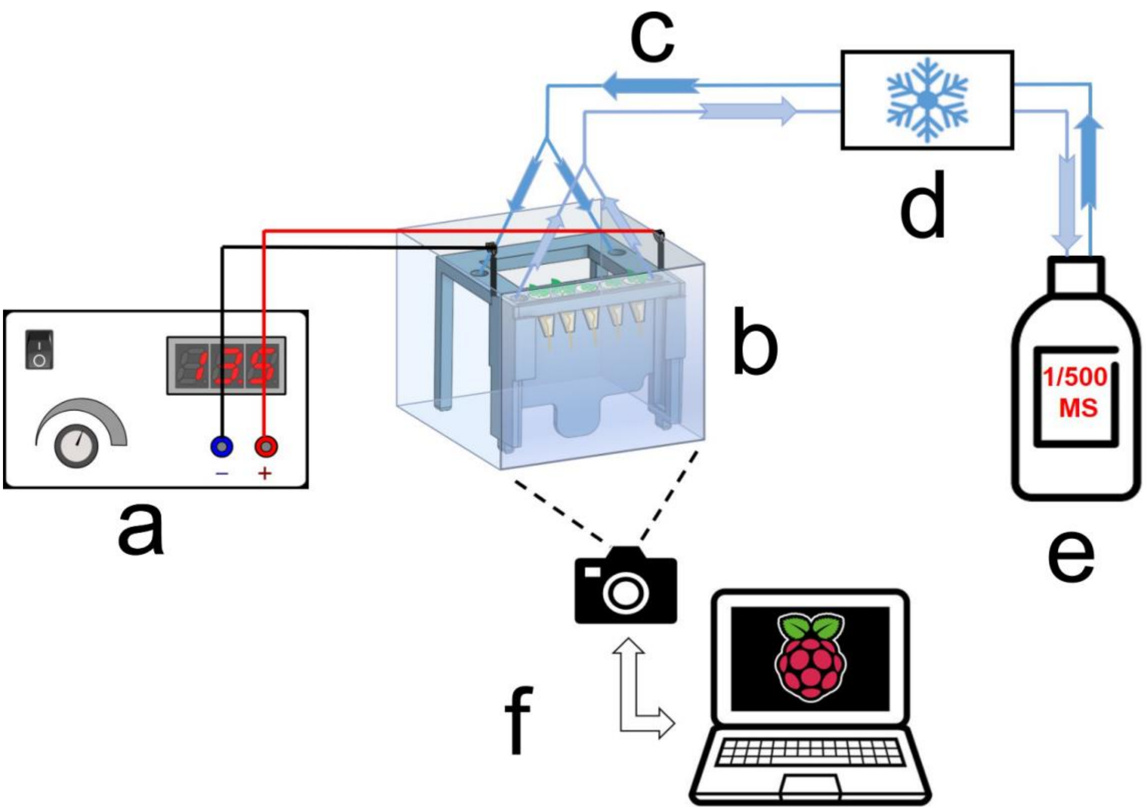


**Supplemental Figure S1.**

Complete electrotropism setup. **(a)** Power supply. **(b)** V-box. **(c)** Flexible tubing for liquid perfusion. **(d)** Chilled water bath. **(e)** Medium reservoir. **(f)** RaspberryPi camera

a)

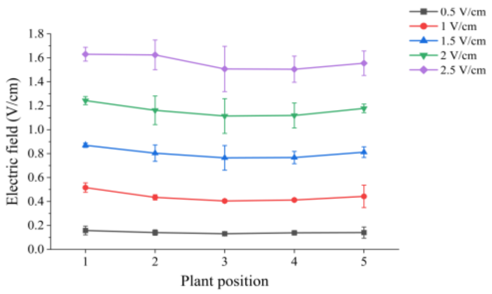


b)

| Electric field (V/cm) | Current (μA) |
|---|---|
| 0.5 | 65.5 ± 5.8 |
| 1 | 157.5 ± 7.9 |
| 1.5 | 247.7 ± 13.3 |
| 2 | 264.7 ± 78.8 |
| 2.5 | 438.5 ± 40.8 |

c)

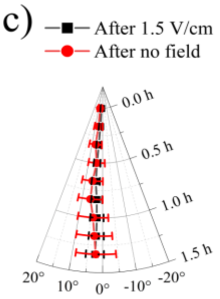


**Supplemental Figure S2**

V-box characterisation. **(a)** Average electric field measurements inside the V-box at each of the five plant positions (N=3 measurements), for each voltage difference imposed by the power supply between the electrodes; error bars, s.d. **(b)** Average (+/- standard deviation) electric currents measured in the circuit for each voltage difference imposed by the power supply between the electrodes. **(c)** Control for efficient circulation of the liquid medium: polar plot of the average root tip orientations with respect to the gravity vector, with time on the radial axis and orientation angle on the circumferential axis, of unexposed roots inside a V-box that had been pre-exposed to 1.5 V/cm or 0 V/cm; error bars, s.e.m.

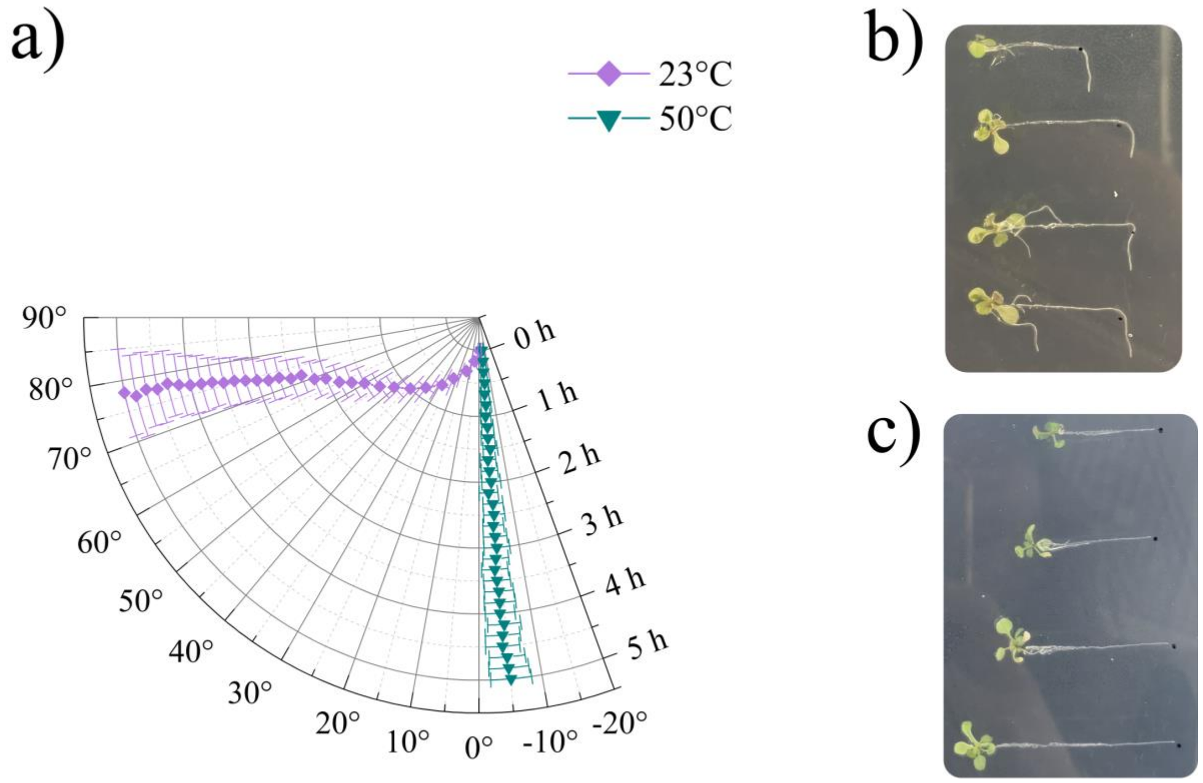


**Supplemental Figure S3**

Live vs dead root. (**a**) Polar plot of the average root tip orientations with respect to the gravity vector, with time on the radial axis and orientation angle on the circumferential axis, for roots pre-treated to 23 ℃ or 50 ℃ and then exposed to 2.0 V/cm; error bars, s.e.m. (**b**) Gravitropism assay after 23 ℃ pre-treatment, representative plants. (**c**) Gravitropism assay after 50 ℃ pre-treatment, representative plants

a)

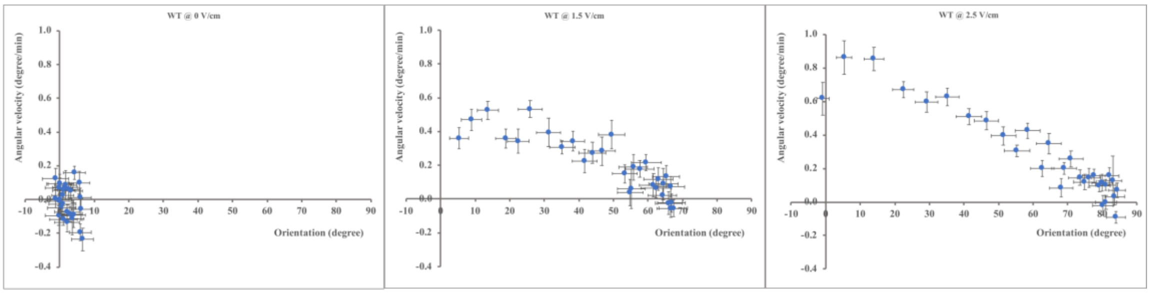


b)

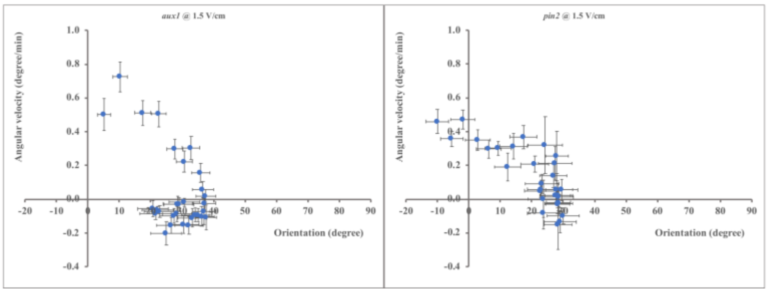


**Supplemental Figure S4**

Velocity of electrotropic response. **(a)** WT root tip average angular velocity, at any observed orientation, for 0 V/cm (left panel), 1.5 V/cm (middle panel) and 2.5 V/cm (right panel); **(b)** same plots for the mutants *aux1* (left panel) and *pin2* (right panel) at 1.5 V/cm. Error bars, s.e.m.

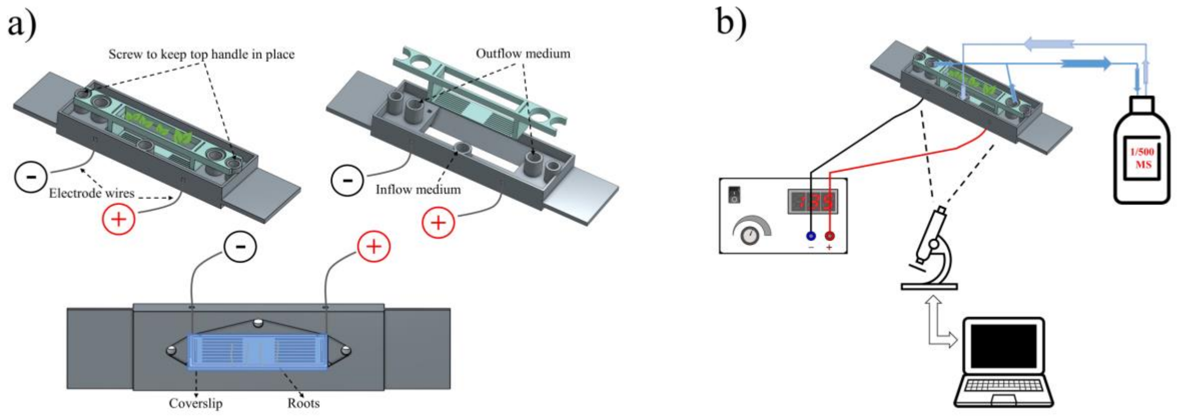


**Supplemental Figure S5**

V-slide. **(a)** Diagram of the V-slide used to image with confocal microscopy a root exposed to the electric field. **(b)** Diagram of the complete setup used with the V-slide.

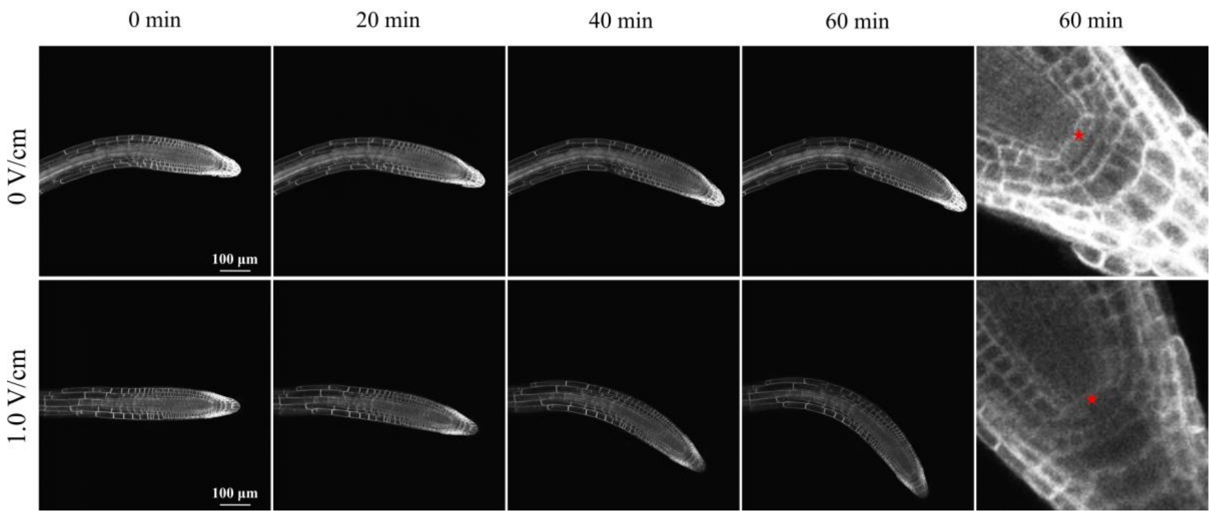


**Supplemental Figure S6**

Root meristem is not damaged. Time-lapse confocal images of two representative roots (total roots observed: N=3 for mock and N=3 for 1.0 V/cm) expressing the WAVE131 marker, exposed to 0 V/cm or 1.0 V/cm in the V-slide; red star, the quiescent centre (QC). Scale bars, 100 µm.

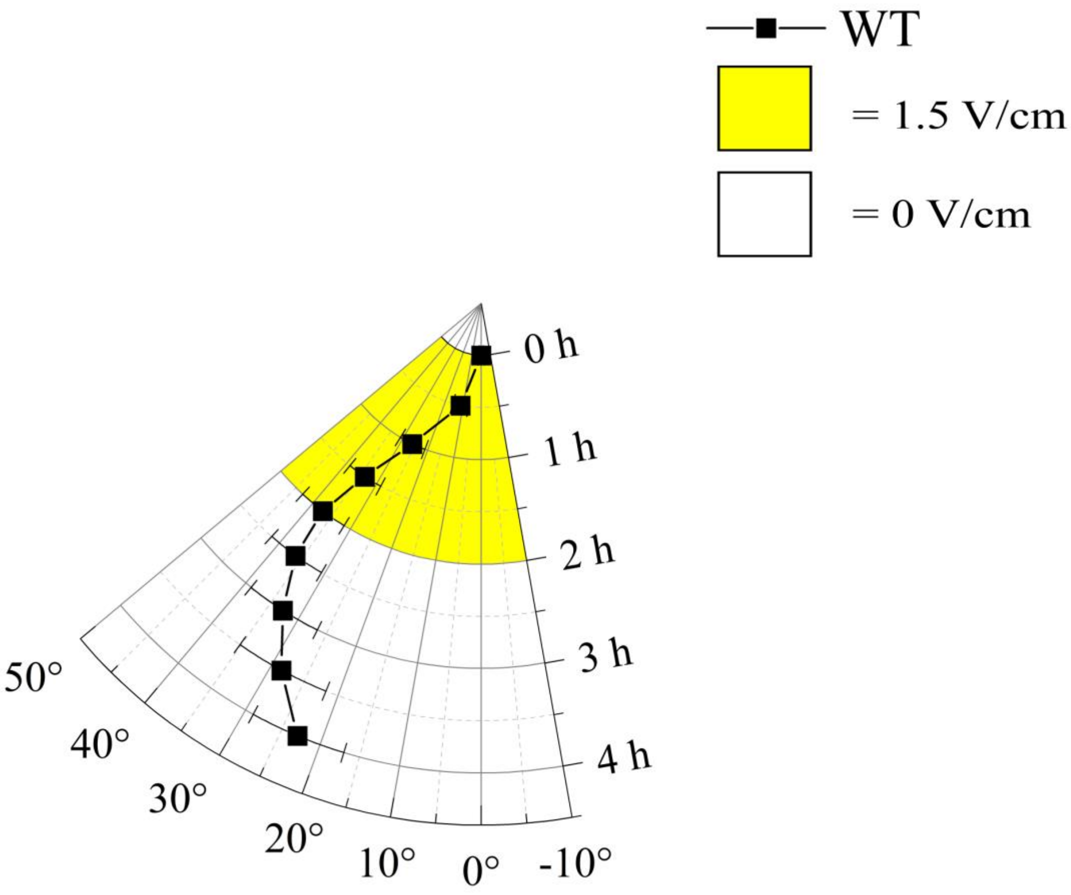


**Supplemental Figure S7**

Roots exposed to the electric field are still gravitropic. Polar plot of the average root tip orientations with respect to the gravity vector, with time on the radial axis and orientation angle on the circumferential axis. Roots have been exposed to 1.5 V/cm for 2 h and then monitored for another 2 h after the field had been turned off. Error bars, s.e.m.
